# SPRAY-INDUCED GENE SILENCING IDENTIFIES PATHOGEN PROCESSES CONTRIBUTING TO POWDERY MILDEW PROLIFERATION

**DOI:** 10.1101/2022.09.13.507682

**Authors:** Amanda G. McRae, Jyoti Taneja, Kathleen Yee, Xinyi Shi, Sajeet Haridas, Kurt LaButti, Vasanth Singan, Igor V. Grigoriev, Mary C. Wildermuth

**Affiliations:** Department of Plant and Microbial Biology, University of California, Berkeley, CA 94720; U.S. Department of Energy Joint Genome Institute, Lawrence Berkeley National Laboratory, Berkeley, CA 94720

**Keywords:** powdery mildew, spray-induced gene silencing (SIGS), dsRNA, *Golovinomyces orontii*, *Arabidopsis thaliana*, *Erysiphe necator*, grapevine

## Abstract

Spray-induced gene silencing (SIGS) is an emerging tool for crop pest protection. It utilizes exogenously applied double stranded RNA to specifically reduce pest target gene expression using endogenous RNA interference machinery. Powdery mildews, widespread obligate biotrophic fungi infect agricultural crops including wheat, barley, cucurbits, grapevine, and ornamentals such as roses. In this study, SIGS methods were developed and optimized for powdery mildews using the known azole-fungicide target *CYP51* and the *Golovinomyces orontii*-*Arabidopsis thaliana* pathosystem. Additional screening resulted in the identification of conserved gene targets and processes important to powdery mildew proliferation: *apoptosis-antagonizing transcription factor* in essential cellular metabolism and stress response; *lipase a, lipase 1*, and *acetyl-CoA oxidase* in lipid catabolism; *9-cis-epoxycarotenoid dioxygenase, xanthoxin dehydrogenase*, and a putative *abscisic acid G-protein coupled receptor* predicted to function in manipulation of the plant hormone abscisic acid; and the secreted effector *EC2*. Powdery mildew is the dominant disease impacting grapes and extensive powdery mildew resistance to applied fungicides has been reported. Therefore, we developed SIGS for the *Erysiphe necator-Vitis vinifera* system and tested six successful targets identified using the *G. orontii-A. thaliana* system. For all targets tested, a similar reduction in powdery mildew disease was observed between systems. This indicates screening of broadly conserved targets in the *G. orontii-A. thaliana* pathosystem identifies targets and processes for the successful control of other powdery mildews. The flexibility, specificity, reduced environmental and health risks, and rapid transition from the bench to the field make SIGS an exciting prospect for commercial powdery mildew control.

## INTRODUCTION

Powdery mildews are widespread obligate biotrophic pathogens that infect a variety of agriculturally important crops (Glawe, 2008). Fungicides are required to control infection, though there are increasing needs for other disease management tools due to increasing prevalence of powdery mildew (PM) resistance to fungicides (Frenkel et al., 2015; Kunova et al., 2021; Miles et al., 2021; Vielba-Fernández et al., 2020) and increasing human health and environmental concerns associated with fungicide and repeated sulfur application (Bastos et al., 2021; Calvert et al., 2008; Hinckley & Matson 2011; Raanan et al., 2017; Tacoli et al., 2020; Zubrod et al., 2014). For example, *Erysiphe necator*, which causes grape PM, is the dominant disease impacting grapes in California and world-wide (Gadoury et al., 2012). Furthermore, PM disease management accounts for 74% of pesticides used by CA grape growers (Sambucci et al., 2014).

RNA interference (RNAi) strategies are being developed to offer additional solutions to combat plant disease. This approach uses conserved eukaryotic host and/or pathogen RNAi machinery to silence the targeted pathogen gene by specifically degrading the targeted messenger RNA, preventing its subsequent translation to protein (Baulcombe, 2004; Majumdar et al., 2017; Zotti et al., 2018). Long double-stranded RNA (dsRNA) specific to the pathogen gene target transcript is processed to multiple small interfering RNAs (siRNAs) of 21 to 28 bp by the endogenous RNAi machinery (Meister & Tuschl, 2004). The siRNA is then incorporated into the RNA-induced silencing complex (RISC) where it guides sequence-specific cleavage of the targeted gene transcript. Initially, RNAi strategies against plant pathogens involved stable introduction of the dsRNA into the host plant, requiring the development of transgenic crops. The resulting host-induced gene silencing (HIGS) was effective in reducing PM (Pliego et al., 2013) and other fungal diseases including *Fusarium* head blight (Machado et al., 2018), wheat stripe rust (Zhu et al., 2017), and gray mold (Xiong et al., 2019). However, despite years of advancement, there are few RNAi-based plant products targeting pests in the marketplace due to plant transformation limitations, the timeline to product development, regulatory burdens, and consumer resistance to transgenic crops (Bramlett et al., 2020). The few crops developed, assessed and approved for human consumption include insect resistant SmartStax corn (Darlington et al., 2022; Head et al., 2017) and virus resistant papaya (Gonsalves, 2006) and plum (Scorza et al., 2013).

Topical application of dsRNA or siRNA, referred to as spray-induced gene silencing (SIGS), offers the possibility of a rapidly tested and field-optimized, non-transgenic and specific approach for limiting crop loss due to pests. Furthermore, the dsRNAs are rapidly degraded in the environment (Bachman et al., 2020), with minimal or no environmental and human health impact. For phytopathogenic fungi, SIGS was first found to be effective in limiting disease of the necrotrophic fungus *Botrytis cinerea* (Wang et al., 2016) and the hemibiotrophic fungus *Fusarium graminearum* (Koch et al., 2016), both Ascomycetes. SIGS was subsequently found to be effective against other fungal pathogens including the obligate biotrophs *Phakopsora pachyrhizi* (Hu et al., 2020), a Basidiomycete, and *Hyalaperonospora arabidopsidis*, an Oomycete (Bilir et al., 2019). However, SIGS is not universally effective against fungi, with efficacy apparently dictated by the efficiency of pathogen RNA uptake (Qiao et al., 2021).

In this study, we establish that powdery mildews, obligate biotrophic Ascomycetes, can readily take up dsRNA, develop an optimized SIGS pipeline for predicting and assessing PM gene target efficacy in reducing PM disease using the *Golovinomyces orontii-Arabidopsis thaliana* system, and translate these methods to the agriculturally relevant system, *Erysiphe necator-Vitis vinifera*. Targets were conserved among PMs and we show 100% translation of our findings with *G. orontii-Arabidopsi*s to *E. necator*-grapevine. Powdery mildews cannot be cultured; thus, it has been difficult to investigate PM gene function. We illustrate how SIGS can be used to interrogate the impact of genes and processes on PM growth and reproduction paving the way for more detailed functional investigations and for RNAi-based control of powdery mildew disease in agriculture and ornamental horticulture.

## RESULTS

### *G. orontii* spores can take up environmental RNA

To determine if *G. orontii* MGH1 *(Gor)* can uptake RNA directly from its environment, germinated conidia were incubated with *in vitro* transcribed Fluorescein-labeled β-actin-Mouse RNA, similar to Wang et al. (2016). Powdery mildew uptake of RNA is rapid and observed within 1.5 hr of application demonstrating PM can uptake RNA independent of the host plant. Fluorescein was observed in all tissues present: germinated conidia, germ tubes, and ungerminated conidia (**Fig. 1**). RNAse treatment decreased background fluorescein signal.

**Fig. 1.**
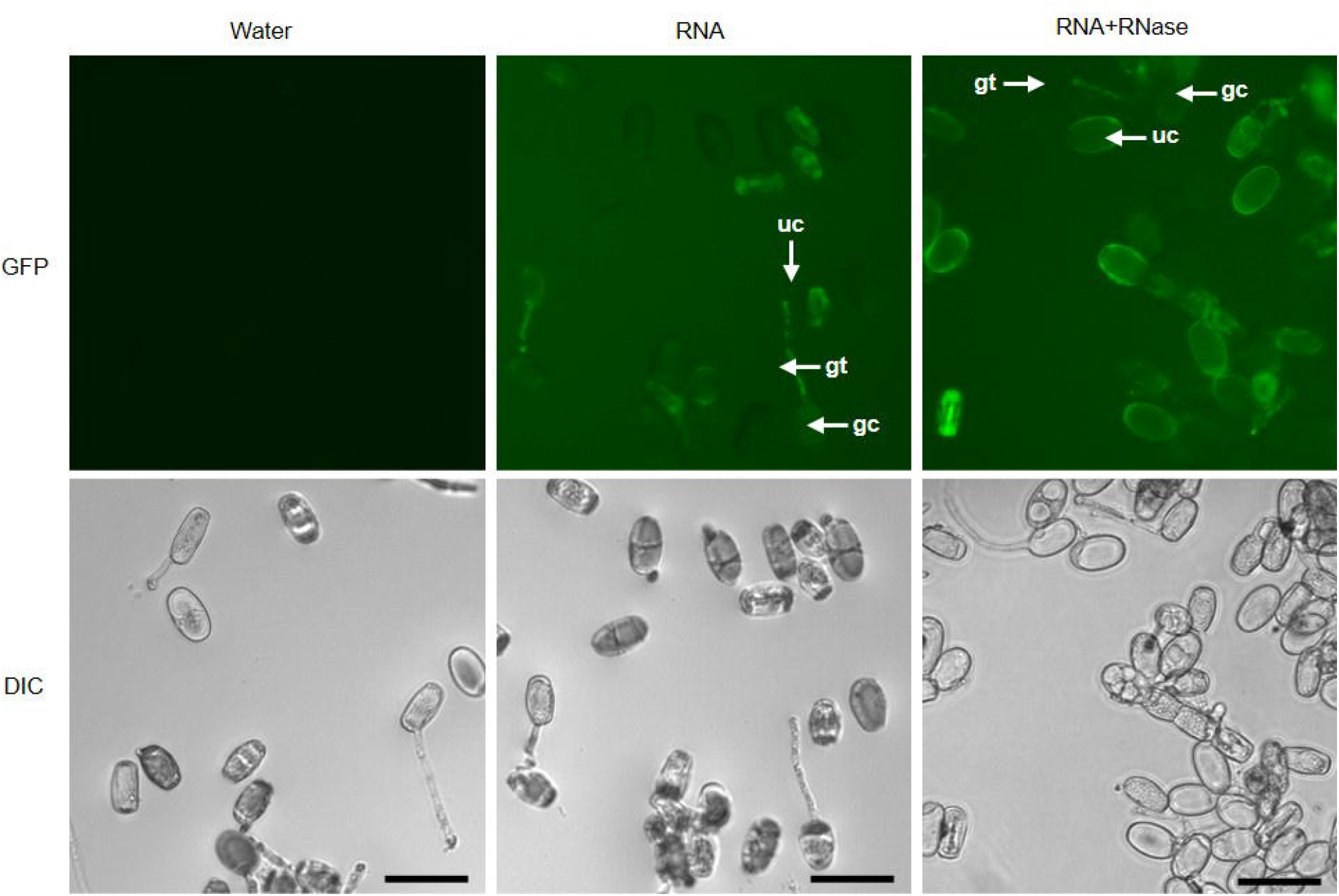
*G. orontii* MGH1 uptakes extracellular RNA. Germinated spores were incubated with water, RNA containing fluorescein-conjugated UTPs (fRNA) or fRNA treated with RNase before imaging. Representative images are displayed; an independent experiment gave similar results. DIC= Differential Interference Contrast. GFP = Green Fluorescent Protein Filter Set. Scale bar= 50 μm. uc= ungerminated conidium; gc= germinated conidium; gt= germ tube.

### siRNA and long dsRNA design

siRNA and/or long dsRNA was designed for each gene target, obtained from published, annotated *Gor* and *E. necator* C-strain (*En*) genomes (Procedures). The online resource pssRNAit was used to identify efficient siRNAs and dsRNAs *in silico* and minimize off-target silencing in the plant host. Long dsRNAs were 199-449 bps and siRNAs were 21-base sense and antisense strands that anneal over 19 bases with 2-base overhangs at the 3’ ends. Target genes and primers are provided in **SI Table 1**.

### siRNA and dsRNA against *GorCYP51* significantly reduces spore production in *A. thaliana* whole plant and detached leaf assays

SIGS methods were optimized using the gene target *CYP51*. CYP51 is a common fungicide target required for the synthesis of sterol, a component of fungal cell membranes (Frenkel et al., 2015). *CYP51* has been successfully silenced using SIGS to reduce growth of the fungal hemibiotrophic pathogen, *F. graminearum* (Koch et al., 2016) and necrotrophic pathogen *B. cinerea* (Nerva et al., 2020). The spraying method, amount of RNA, number of RNA applications, timing of RNA application, RNA purification methods, and detached leaf and whole plant protocols were optimized to improve efficiency and reproducibility of results. **Figure 2** provides an overview of our SIGS screening protocol with methods described in Procedures.

**Fig. 2.**
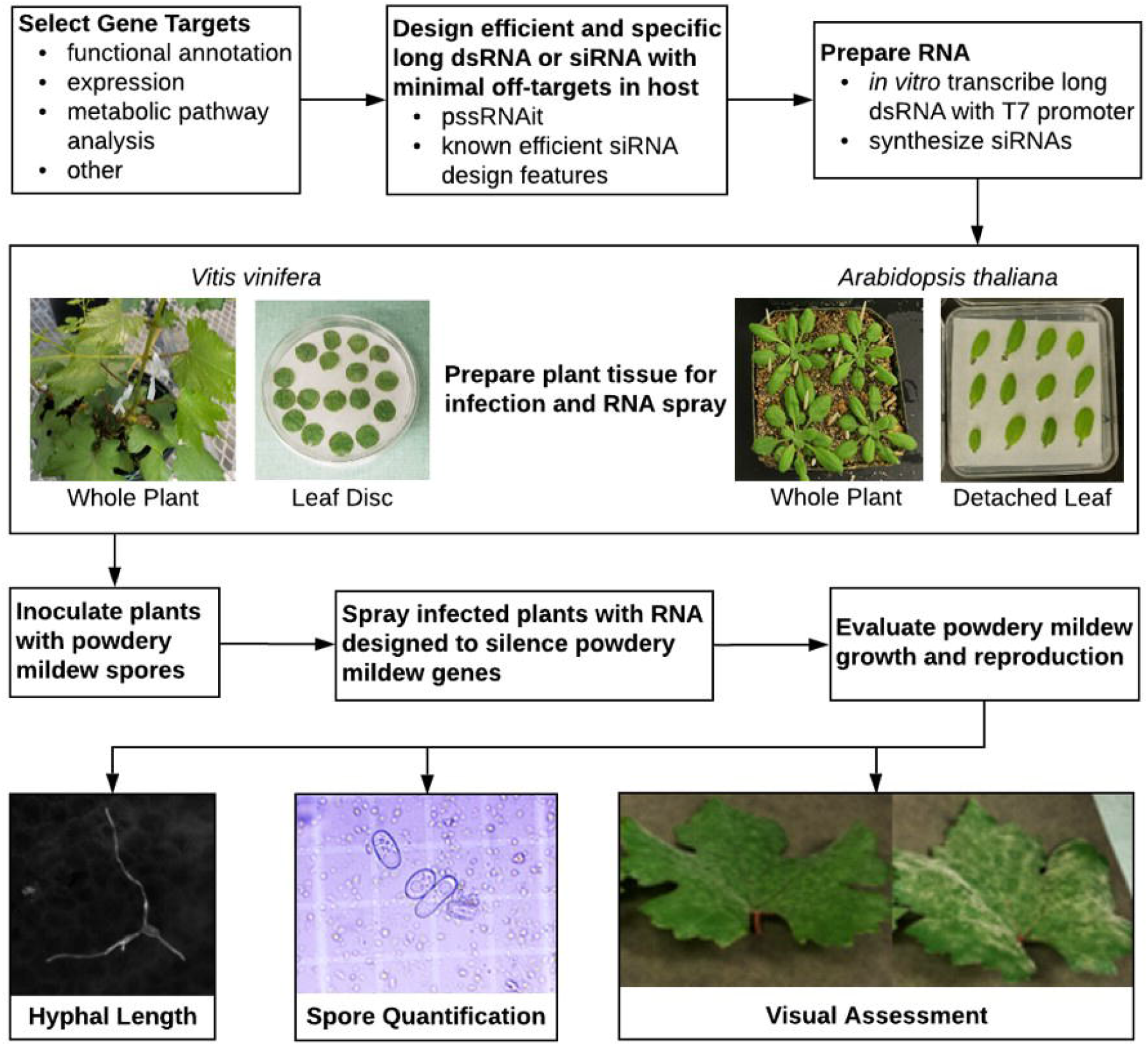
Flowchart of spray-induced gene silencing methodology.

Two 21 bp siRNAs, siRNA-1 and siRNA-2, and two long dsRNAs, dsRNA-1 and dsRNA-2, targeting *GorCYP51* were designed (**Fig. 3a, SI Table 1**). In whole plant and detached leaf experiments using *CYP51* dsRNA-1, spore production at 8-10 days post inoculation (dpi), decreased by an average of 46% and 30% respectively (**Fig. 3b**) and less PM coverage and density was observed on dsRNA-sprayed plants than buffer control-sprayed plants (**Fig. 3c**). Surprisingly, dsRNA-2 was less reproducible in limiting PM spore production despite significant overlap with dsRNA-1. However, siRNA-1 reproducibly reduced spore production by 35% and 40% in whole plant and detached leaves, respectively, while siRNA-2, which targets a different region of *CYP51*, did not significantly reduce powdery mildew (**Fig. 3b**).

**Fig. 3.**
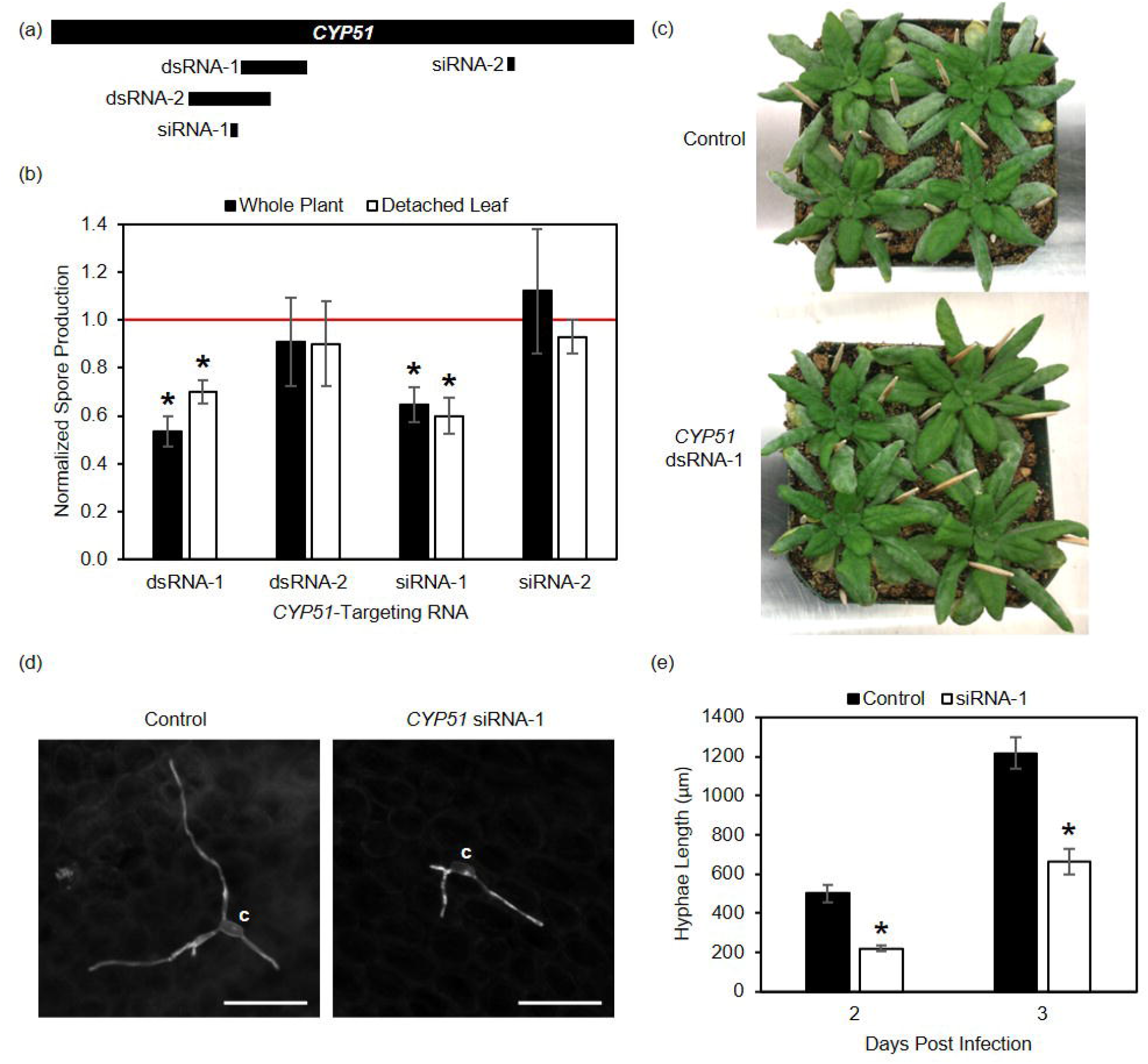
siRNA and long dsRNA targeting *CYP51* reduces spore production and hyphal length. (a) Diagram of *CYP51* transcript (1578 bp) and designed inhibitory RNAs. siRNAs were synthesized. siRNA-1:GUACGUGCCCAUAAUUCAAAA/UUGAAUUAUGGGCACGUACGA. siRNA-2: CGGAUGUACUAGUCGUGAUCC/AUCACGACUAGUACAUCCGGG. See SI Table 1 for long dsRNA primers. (b) Mean spore production normalized to control parallel samples; n≥3 (c) Powdery mildew on representative control and parallel *CYP51* dsRNA-1 sprayed *A. thaliana* plants, at 9 dpi. (d) Representative images of calcofluor-stained *G. orontii* on control and siRNA-1 treated leaves. Scale bar= 100 μm; c= conidium. (e) Mean hyphal length per PM colony from detached leaf experiments; n≥35. Error bars ± SEM. *p<0.05 by Student’s T-Test (unpaired, 2-sided) compared to control. Similar results observed in independent experiments.

These results show SIGS against *GorCYP51* can reproducibly limit PM disease of *Arabidopsis*. However, the efficacy is dependent on the exact dsRNA or siRNA employed. This indicates a negative result does not mean the targeted transcript is not important to the phenotype assessed; instead, the negative result can be due to poor silencing efficiency of a particular siRNA or long dsRNA. The success of siRNA-1 but not dsRNA-2 (within which siRNA-1 resides) treatment in limiting spore production suggests that differential dsRNA uptake and/or processing of dsRNA-2 to siRNAs may be important. Finally, the fact that not all dsRNA or siRNA tested alter *Gor* spore production indicates that it is not the addition of dsRNA or siRNA itself that impacts PM growth and reproduction.

### Targeting *CYP51* significantly reduces *Gor* hyphal length on *A. thaliana* leaves

To see the impact of SIGS on earlier PM colony development, the impact of *CYP51* siRNA-1 on hyphal length was measured at 2 and 3 dpi in *A. thaliana* detached leaf assays. At this stage, nutrients are taken up by the fungus through the haustoria and the colonies are continuing to expand but asexual reproduction is not yet occuring. The hyphal branching architecture was not altered (**Fig. 3d**); however, the average total hyphal length of *CYP51* siRNA-1 colonies was significantly reduced at 2 and 3 dpi, by 56% and 45% respectively, compared to control samples (**Fig. 3e**). The reduction in both early *Gor* hyphal growth and spore production with *CYP51* siRNA-1 treatment is consistent with the essential role of CYP51 in fungal membranes and its consistent expression through all stages of infection (**Fig. 4**).

**Fig. 4.**
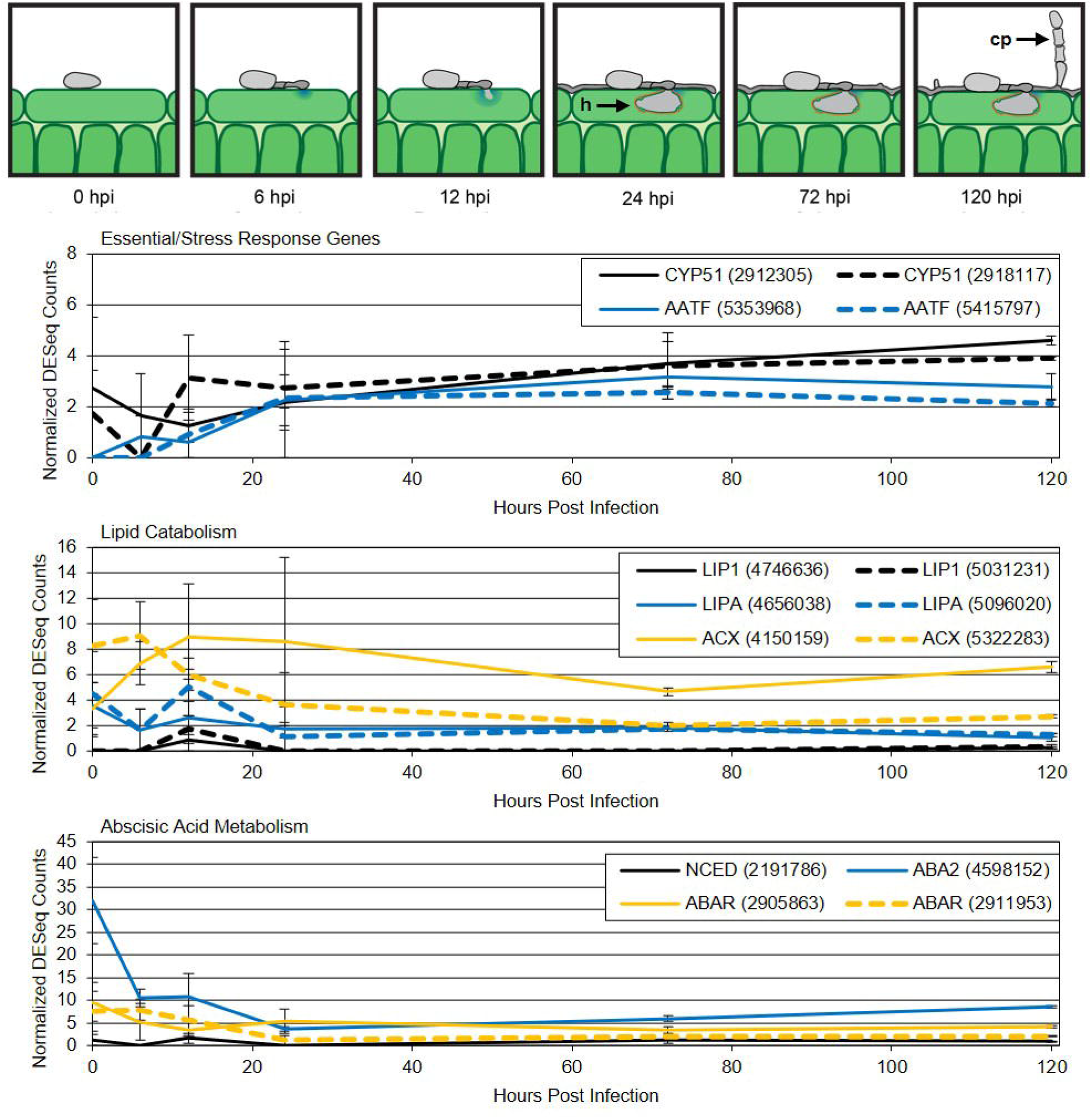
Time course of *G. orontii* development on *Arabidopsis* with *G. orontii* MGH1 gene expression shown for SIGS PM targets by function. Developmental timeline: inoculation (0), spore germination (6), appressorium development and cell wall penetration (12), haustorial maturation (24), colony expansion (72), and asexual reproduction (120) hrs post inocuolation (hpi). Mean DESeq2 normalized gene expression (± SEM) from triplicate samples. Most *Gor* gene targets have two copies due to recent duplication. h= haustoria; cp= conidiophore, asexual reproductive structure with new conidia.

### PM gene selection for SIGS analysis

For subsequent SIGS analysis, we prioritized broadly conserved PM genes with the following predicted functional roles: 1. essential cellular function and stress response, 2. energy production, and 3. manipulation of the plant host. Genes involved in essential cellular functions, such as *CYP51*, are expected to be effective targets as they have been for chemical fungicides. In addition, genes that moderate cellular stress can be essential to growth and reproduction of the PM on its host. Although PMs evade activating plant defense and limit those defenses that do occur, they are exposed to damaging compounds such as reactive oxygen species and specialized metabolites produced at the infection site. Energy production is critical to PMs as it is for all organisms. During early colonization, the PM derives its energy from the breakdown of energy-rich compounds in the spore. For example, PM spores are replete with glycogen and triacylglycerols (TAGs) that are mobilized and degraded during germination and appressorial development (Both et al., 2005). Once the fungal feeding structure (haustorium) has developed, PMs rely on plant resources as they have lost the ability to synthesize costly compounds readily obtained from the host plant (Spanu, 2012). Many plant pathogens including PMs manipulate plant functions such as cell fate, phytohormone metabolism, and defense through the secretion of effector proteins that enter the plant cell (Lo Presti et al., 2015; Weßling et al., 2014). Furthermore, plant pathogens can directly manipulate plant hormone responses by making a plant hormone or hormone (ant)agonist (Ronald & Joe, 2018).

When selecting specific genes from the above functional categories for prioritization, conservation in PMs, functional annotation, metabolic pathway, and pattern of expression were taken into consideration. Metabolic pathway analysis included annotation and pathway mapping using KEGG (Kanehisha & Goto, 2000), and prioritization of rate-limiting enzymes that dominate flux through a pathway and/or most highly regulated enzymes in a pathway. *Gor* gene expression over the course of *A. thaliana* infection employed RNASeq data at 0 (inoculation), 6 (spore germination), 12 (appressorial development and plant cell wall penetration), 24 (haustorium), 72 (continuted colony expansion - hyphal growth and formation of secondary haustoria), and 120 (asexual reproduction) hrs post inoculation. PM gene targets involved in essential cellular function and stress response, *apoptosis-antagonizing transcription factor* (*AATF)*, and in energy production from glycogen (*glycogen debranching enzyme 1* (*GDB1*), *glycogen phosphatase 1* (*GPH1*), *glucose-induced degradation 9* (*GID9*)) and lipids (*lipase 1* (*LIP1*), *lipase A* (*LIPA*), *acyl-coenzyme A oxidase* (*ACX*)) were selected. In addition, a putative *diglyceride acyltransferase* (*DGAT*) catalyzing the formation of TAGs for energy storage was examined. PM targets *9-cis-epoxycarotenoid dioxygenase* (*NCED*), the xanthoxin dehydrogenase *absicic acid 2 (ABA2)*, and the putative *absicic acid G-protein coupled receptor (ABAR)* are predicted to function in the manipulation of the plant host via the plant hormone abscisic acid (ABA). Finally, the highly expressed secreted protein *effector candidate 2* (*EC2*) was targeted. We screened multiple genes known or predicted to be involved in functional categories of interest to increase our probability of identifying functional categories of highest importance to the powdery mildew. These genes tended to have similar levels and patterns of expression (**Fig. 4, SI Fig. 1**). The twelve *Gor* gene targets comprise 8 metabolic genes, 3 regulatory genes, and 1 gene encoding a secreted effector protein and show different levels and patterns of expression over the course of infection.

### dsRNA designed to target metabolic, regulatory, and effector PM genes identifies targets and processes that contribute to *Gor* spore production on *A. thaliana*

Using our optimized SIGS protocol (**Fig. 2**) with long dsRNA and the spore production assay, we tested twelve new PM targets. Spore production integrates all facets of PM development, is most relevant to growers, and is faster than microscopic assays which assess earlier phases of PM development such as hyphal growth. We found SIGS against 8 of 12 new targets tested significantly reduced PM proliferation (**Fig. 5**). The 8 targets with reduced PM spore production on whole plants were also assessed using the detached leaf assay; SIGS against all 8 resulted in significantly reduced PM spore production. While SIGS against all three PM targets known or predicted to function in lipid catabolism (*LIP1, LIPA, ACX*) or manipulation of the plant hormone ABA (*NCED, ABA2, ABAR*) reduced PM proliferation, none of the three targets (*GDB1,GPH1, GID9*) involved in glycogen metabolism were effective. In addition, SIGS against *AATF* and the effector *EC2* reduced spore production; however, SIGS against the putative *DGAT* did not, either due to functional redundancy as other genes encoding enzymes with putative DGAT activity are present in *Gor* or because the dsRNA tested was not efficient at silencing.

**Fig. 5.**
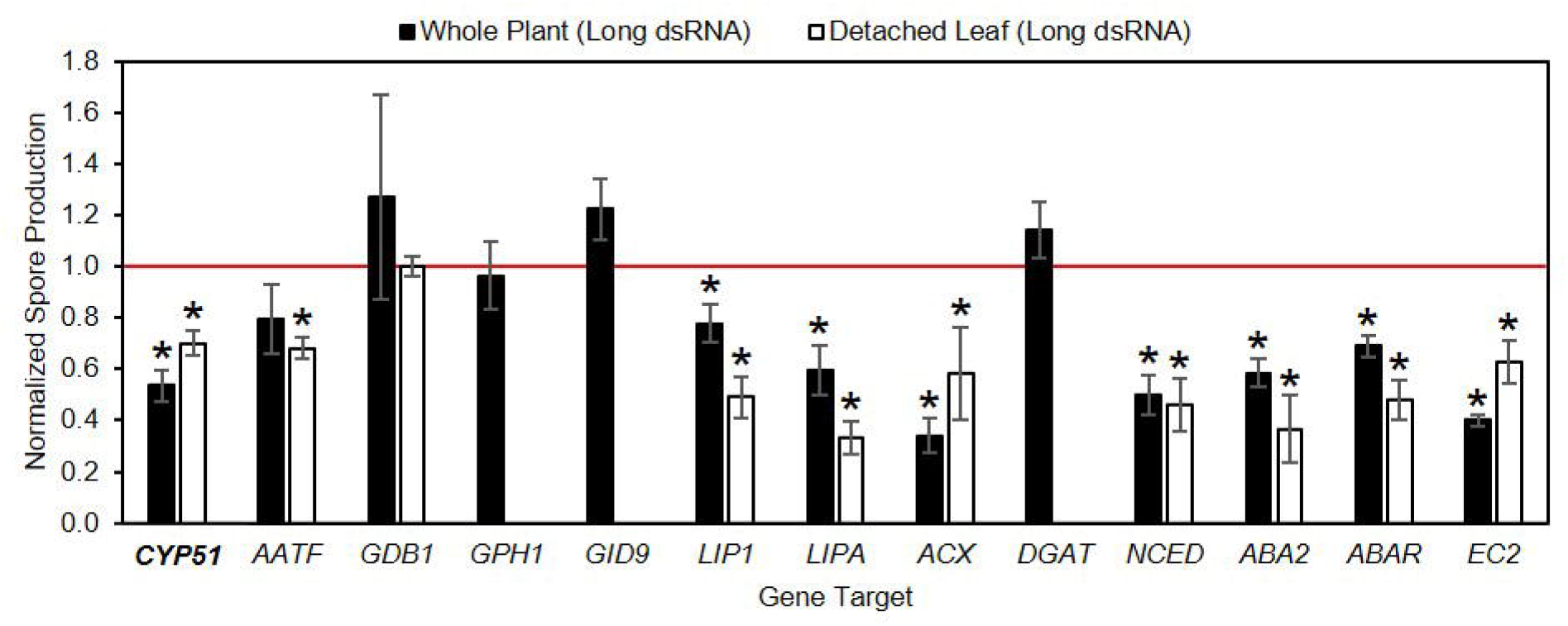
*G. orontii* MGH1 spore production on *A. thaliana* is impacted by long dsRNA targeting diverse PM genes. Mean normalized spore production ±SEM displayed from n ≥ 3. *p<0.05 by Student’s T-Test (unpaired, two-tailed) compared to control. Independent experiments gave similar results.

### SIGS methods and gene targets translate to the *Erysiphe necator-Vitis vinifera* pathosystem

We next sought to translate these methods and targets to grape powdery mildew (**Fig. 2**). New long dsRNAs were designed for each gene tested as there is sequence divergence between *Gor* and *En* transcripts; however, the same region of the gene was targeted. First, procedures were optimized for the grapevine system with *EnCYP51* as the gene target. Spore production was reduced by 62% and 45% in grapevine whole plant and leaf disc assays, respectively, when treated with long dsRNA against the same region of *EnCYP51* as dsRNA-1 against *GorCYP51* (**Fig. 6a**). In addition, visual inspection showed infected grape leaves had less PM coverage and density when sprayed with dsRNA targeting *EnCYP51* than with water alone (**Fig. 6b**). We then tested SIGS against *En AATF, LIP1, LIPA, NCED*, and *EC2* in the grapevine system. SIGS using dsRNA against these genes reduced *En* spore production 53-64% (**Fig. 6a**).

**Fig. 6.**
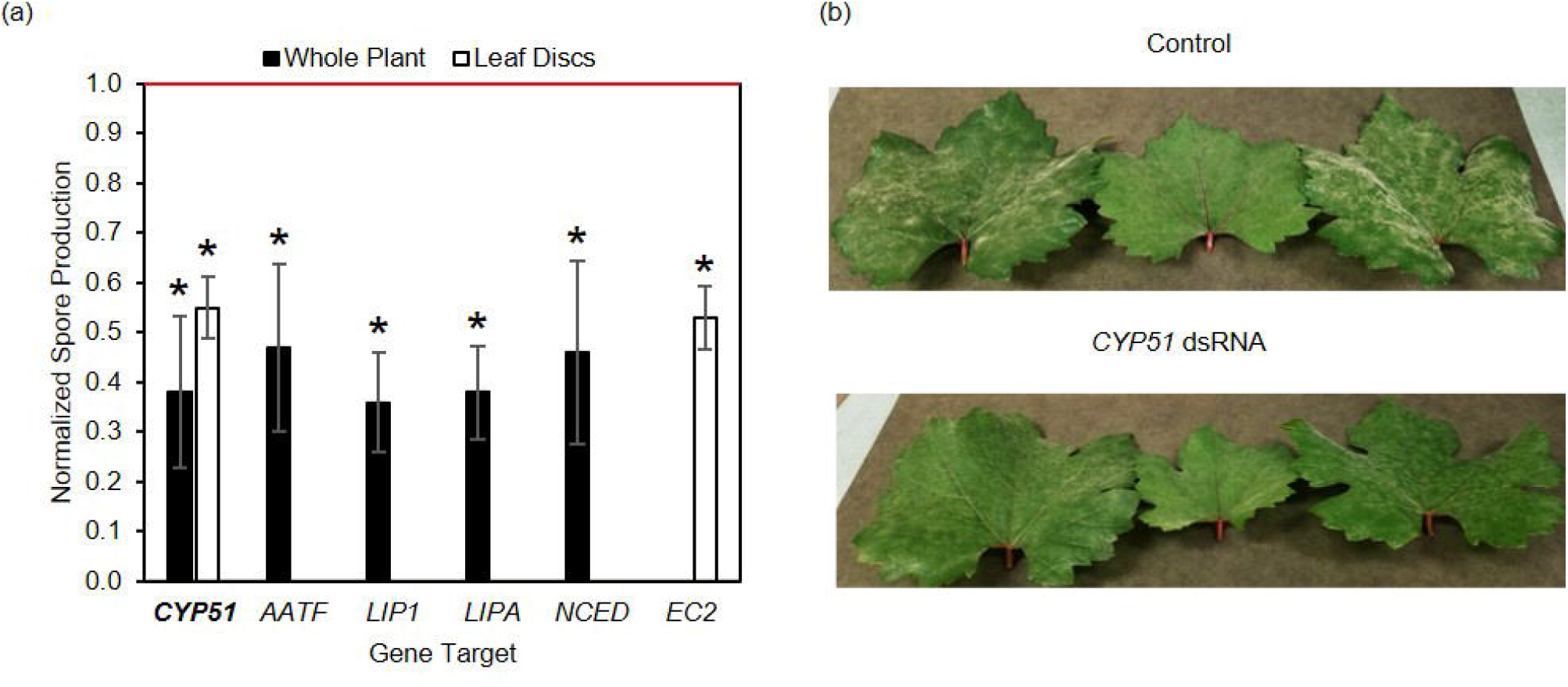
Long dsRNA targeting genes in *E. necator* reduces spore production on *V. vinifera*. (a) Mean normalized spore production ± SEM, n ≥ 3. Note n=2 for AATF. *p<0.05 by Student’s T-Test (unpaired, two-tailed) compared to control. (b) Powdery mildew on control and *CYP51* dsRNA-treated grapevine leaves at 16 dpi.

In summary, six diverse PM genes for which SIGS effectively limited spore production of *Gor* on *Arabidopsis* were tested against the orthologous *En* genes in grapevine, and all six showed a similar reduction in PM growth and reproduction using the grapevine system. They represent targets with known or predicted functional roles in essential cellular function and stress response (*CYP51, AATF*), lipid catabolism (*LIP1, LIPA*), and manipulation of the plant host via ABA hormone metabolism *(NCED*) or effector *(EC2*). This gives us a 100% success rate in translation of our findings from the *Gor*-*Arabidopsis* system to the *En*-grapevine system. It further illustrates the power of using SIGS with the more rapidly assessed *PM-Arabidopsis* system to identify conserved PM targets and processes critical to PM growth and reproduction.

## DISCUSSION

Herein, we develop a pipeline to silence powdery mildew genes and determine their contribution to disease (**Fig. 2**). Screening PM gene targets in *Arabidopsis* is rapid and provides insight into genes important to diverse PMs, as ∼75% of all PM genes are conserved (Wu et al., 2018). *A. thaliana* plants have a quick generation time, grow well in environmentally controlled growth chambers, and require little space for multiple replicates, compared to most PM-impacted crops, making this system valuable for the rapid identification of effective SIGS targets. We also demonstrate our SIGS methodology and effective targets in *Gor* translate to the grape PM *En*, and we anticipate translation to other powdery mildews of importance to agriculture and ornamental horticulture.

Because powdery mildews are obligate biotrophs that cannot be cultured, it has been difficult to interrogate the function of PM proteins. Recently, Ruiz-Jimenez et al. (2021) showed SIGS against the melon PM *Podosphaera xanthii* can be used to identify conserved proteins with no annotated functional domain that are important to PM proliferation; SIGS against three such targets dramatically reduced PM coverage on melon leaves. We show SIGS can rapidly identify genes and processes that contribute to PM growth and reproduction. Although we chiefly employ an endpoint assay of spore production in these analyses, earlier stage-specific impacts can also be assessed (**Fig. 4**). One such assessment, hyphal growth, is shown in **Figure 3**. Examination of the pattern of expression for PM genes of interest can provide indicators of potential phase-specific functions. For example, CYP51 is required for the synthesis of an essential fungal cell wall sterol and thus anticipated to be required throughout infection and growth. Indeed, its expression is constant **(Fig. 4**), and SIGS targeting *CYP51* decreases both early hyphal growth (assessed at 2-3 dpi) and spore production (assessed at 8-10 dpi) by similar amounts (**Fig. 3**).

Using SIGS we probed the importance of thirteen powdery mildew genes in total and identified nine PM targets that significantly impact *Gor* spore production (**SI Table 1, Fig. 5**). These nine PM targets represent metabolic, regulatory, and secreted effector protein categories proposed or known to function in essential cellular function and stress response, energy production, or manipulation of the plant host. Furthermore, the efficacy of SIGS was independent of the level of target gene expression or the pattern of expression (**Figs. 4, 5**). The nine gene targets with demonstrated importance to PM growth and reproduction include two sets of three gene targets predicted to function in the same processes: lipid catabolism and manipulation of the plant hormone abscisic acid. Furthermore, all three glycogen metabolism targets show no SIGS phenotype.

This highlights the utility of screening multiple genes predicted to function in the same process. If SIGS against one of the three targets in a given functional process does not impact PM spore production, it would suggest further dsRNAs against that target be tested as not all long dsRNAs or siRNAs selected using current guidelines are equally effective in silencing a given target (e.g. **Fig. 3**). Similarly, once a phenotype has been observed with SIGS against a given target, further optimization of the applied RNA sequence may further increase efficacy. The efficiency of dsRNA-mediated silencing depends on several factors including uptake, processing to siRNAs, accessibility of the target region of the gene, and structural features of the siRNA resulting in efficient loading of the siRNA antisense strand into the RISC complex (Fakhr et al., 2016; Reynolds et al., 2004). Understanding features of gene hot spots for effective RNA silencing is an active area of investigation, and as discovered, these features will further inform our RNA design. In the meantime, we demonstrate how assessing SIGS against multiple targets known or predicted to act in the same functional process can identify those that should be prioritized for additional effort. Below, we further discuss the successful PM SIGS targets and their predicted functions in the powdery mildew-host interaction.

### Essential cellular function and stress response

Similar to the essential sterol demethylase *CYP51*, the PM *apoptosis antagonizing transcription factor AATF* exhibits low-level expression throughout development including asexual reproduction on the leaf (**Fig. 4**). SIGS against *AATF r*educes PM spore production on Arabidopsis, with an even greater impact (45% reduction) on *En* infection of grapevine (**Figs. 5, 6**). AATF (also known as Che-1) is a eukaryotic cell-fate transcriptional regulator known to suppress apoptosis, promote proliferation (and tumors), mediate the DNA damage response, and integrate cellular stress response (Iezzi & Fanciulli, 2015; Kaiser et al., 2020). Most studies have been done in animal systems, and AATF function in fungi has not been explicitly explored. In fungi, regulated cell death is a factor in fungal development, response to external compounds/stress, and heterokaryon incompatibility (Gonçalves et al., 2017). Although inhibition of plant cell death is critical to PM growth and development, little is known about the role of regulated cell death in obligate biotrophic fungi, including PMs. MycoCosm analyses (Grigoriev et al., 2014) found AATF orthologs with the same domain structure to be present in all high-quality PM genomes and a small subset of other *Leotiomycetes* (Procedures). Given the potentially critical role for AATF in PM stress response and development, optimization of the SIGS dsRNA and/or siRNA targeting AATF is warranted to further increase efficacy. And, other targets that mediate cell fate, and in particular apoptosis and DNA repair in response to genotoxic stress (including exposure to specific specialized plant metabolites), should be explored.

### Lipid catabolism

Energy-rich storage lipids are highly abundant in the PM spore and have been shown to be catabolized and/or mobilized during spore germination, appressorium formation and penetration before the haustorium has fully developed (Both et al., 2005). After haustorium formation, the PM can acquire plant storage lipids (TAGs and/or their breakdown products) via this interface to fuel its continued growth and reproduction. At asexual reproduction, new PM spores filled with storage lipids develop further increasing fungal demand for these precursors. Furthermore, lipids can function as signaling molecules in plants and fungi (Siebers et al., 2016). The three PM gene targets *LIP1, LIPA*, and *ACX* predicted to function in lipid catabolism exhibit low level expression throughout the infection process with *LIP1* and *LIPA* exhibiting slightly elevated expression at 12 hpi, before the haustorium has fully formed (**Fig. 4**). SIGS against each of these targets reduces spore production by >50% providing direct experimental evidence of their importance to PM growth and development.

Lipase 1 contains a secretion signal and a carboxylesterase family protein domain; it is predicted to catalyze the hydrolysis of TAGs into diacylglycerols and a carboxylate. *Blumeria graminis* f. sp. *tritici* (*Bgt*) Lip1 localizes to the surface of conidia, germ tubes, and/or hyphae, and the enzyme has broad lipid specificity with the capacity to hydrolyze leaf epicuticular waxes (Feng et al., 2009). This function of *Bgt*Lip1 has been associated with promoting PM early colonization events via enhanced adhesion to the leaf surface and/or enhanced rate of appressorial germ tube formation (Feng et al., 2009); however, loss of function has not been explored. *LIP1* mutants of the plant necrotroph *B. cinerea* and hemibiotroph *F. graminearum* show no defects in colonization or growth (Subramoni et al., 2010). This may be attributed to the plethora of *LIP1*-like genes in these fungi, compared with obligate plant biotrophs (Feng et al., 2009), minimizing the impact of single *LIP1* mutants. Alternatively, the function of Lip1 may be more critical for the obligate biotrophic PMs. For example, Lip1 activity may release specific cues from the plant host surface that promote PM development (Feng et al., 2009) or provide energy sources for PM growth and reproduction. Here, using SIGS against *LIP1* in the *Gor-Arabidopsis* and *En-grapevine* systems, we directly show the important role that Lip1 plays in PM growth and development with >50% reduction in spore production (**Figs. 5, 6**).

Lipase A contains a secretion signal and a class 3 lipase domain (PF01764; Mistry et al., 2021); as such, it is predicted to function as an extracellular TAG lipase. Orthologs of LipA are present in both plant and animal fungal pathogens, and we found *LIPA* to be present in all PMs and *Leotiomycetes* analyzed including the plant necrotrophs *B. cinerea* and *S. sclerotiorum*. FGL2 (fg01240;ABW74155.1), the *F. graminearum LIPA* ortholog, has lipolytic activity and plays a critical role in virulence on both maize cobs and wheat heads (Nguyen et al., 2008, 2010). We found SIGS against *LIPA* results in a dramatic reduction in PM spore production for both *Gor* (67%) and *En* (62%) (**Figs. 5, 6**). Operating in the extracellular space, GorLipA could act on plant TAGs to release free fatty acids for fungal acquisition. In addition, the released free fatty acids could function as signaling molecules.

SIGS against *GorACX*, peroxisomal acetyl-CoA oxidase (ACX), was also highly effective, reducing spore production by 66% in whole plant assays (**Fig. 5**). ACX participates as a rate-limiting enzyme in fatty acid B-oxidation and □-linolenic acid metabolism. Therefore, in PMs, ACX could be involved in degradation of long chain free fatty acids for energy generation or the production of oxylipins as fungal reproductive signals or as plant hormone jasmonic acid (ant)agonists (Poirier et al., 2006).

### Abscisic acid metabolism

The plant hormone ABA plays a critical role in diverse plant processes including seed development, dormancy, and germination; regulation of growth and development; stomatal closure; and response to abiotic and biotic stressors (Jia et al., 2022). Fungal production of ABA appears to be limited to plant-associated fungi, acting as a virulence factor to mediate ABA-dependent plant responses (Takino et al., 2019). Fungi and plants employ distinct ABA biosynthetic routes (Jia et al., 2021; Takino et al., 2019). However, the annotated *Gor* genome contains two plant ABA biosynthetic enzymes, NCED and ABA2, and reciprocal protein BLAST analysis found the *Gor* proteins to be the best protein BLAST hits of the Arabidopsis plant proteins and *vice versa* (Procedures). By contrast, no *Gor* ortholog of the fungal terpene synthase (e.g. BcABA3 in *Botrytis cinerea*) that catalyzes the first step of fungal ABA synthesis was identified. NCED (EC 1.1.3.11.51), the rate-limiting enzyme in plant ABA synthesis, cleaves 9-cis violaxanthin and 9-cis-neoxanthin to xanthoxin. ABA2 (EC 1.1.1. 288), a xanthoxin dehydrogenase, then converts xanthoxin to abscisic aldehyde. The specificity of the enzyme functions and their sequential activities suggest fungal acquisition of these genes from the plant, as does the absence of *NCED* in PMs of monocots (Wisecaver & Rokas, 2015). However, it should be noted that *GorNCED* resides in a gene cluster involved in fungal carotenoid metabolism and fungi are capable of making a diverse array of carotenoid-derived products with functions in signaling, development, and environmental response (Avalos & Limon, 2015).

While *Gor* does not contain a plant PYR/RCAR ABA receptor (Ma et al., 2009: Park et al., 2009) homolog, a putative plant ABA G-protein coupled receptor (ABAR) (Pandey et al., 2009) homolog was identified in *Gor*. GorABAR is a nine-transmembrane domain protein with an N-terminal GPHR domain (PF12537) and an ABA GPCR domain (PF12430) (Mistry et al., 2021). The *Gor* and Arabidopsis ABAR proteins are the reciprocal best protein BLAST hits of each other. Furthermore, all PMs contain a highly conserved putative ABAR, while other *Leotiomyctes* analyzed did not (Procedures). While the PYR/RCAR ABA receptors account for dominant ABA-associated plant responses, the Arabidopsis ABA GPCRs can bind ABA *in vitro* (Pandey et al., 2009) and thus could potentially function as an ABA or ABA intermediate receptor in fungi.

The expression patterns of *Gor NCED, ABA2*, and *ABAR* are fairly similar with peaks at 0 and/or 12 hpi and some expression throughout (**Fig. 4**), with *ABA2* exhibiting the highest level of expression and a more dramatic peak of expression (at 0 hpi). SIGS against each of these genes led to a dramatic and similar decrease in PM spore production of 63 to 52%. This pattern of expression and function could be consistent with a known role for ABA - impacting fungal appressorium formation and penetration. For the hemibiotrophic rice pathogen, *Magnaporthe oryzae*, exogenous application of ABA promotes fungal spore germination and appressoria formation (Spence et al., 2015). Furthermore, a *M. oryzae* ABA biosynthetic mutant exhibits reduced appressoria formation and lesion formation on rice. Powdery mildew infection alters the expression of ABA biosynthetic and responsive genes in the host plant (Chandran et al., 2010; Hayes et al., 2010). This could be associated with ABA promotion of the plant source-to-sink transition associated with PM infection, alteration of plant defenses, and/or localized drought response. A recently identified PM effector targets the plant ABA biosynthetic pathway (Li et al,. 2020) confirming the importance of its regulation. While the role of host ABA in PM infection and growth needs to be further resolved, our findings introduce additional complexity to the interaction as the PM itself may be able to synthesize ABA intermediates to manipulate host function, or for its own, yet undetermined, purposes.

### PM effector manipulation of the plant host

Effectors are secreted microbial proteins that enter the plant cell to manipulate plant functions. The effector EC2 is a widely conserved and highly expressed PM effector first described in *B. graminis* f.sp. *hordei (Bgh)* as BEC2 (CSEP0214; Schmidt et al., 2014). EC2 contains a cysteine-rich fungal extracellular membrane (CFEM) domain that is unique in fungi and contains eight conserved cysteine residues. Fungal CFEM-containing proteins have been shown to play roles in fungal pathogenesis and development (Kou et al., 2017; Zhu et al., 2017). We found PM EC2 effectors form their own cluster within fungal CFEM-containing protein domain architectures (Procedures). *GorEC2* is extremely highly expressed, at levels 1-2 orders of magnitude higher than the other genes targeted in this study (**Fig. 4)** as are other PM *EC2* orthologs (e.g., Fonesca et al., 2019; Jones et al., 2014; Schmidt et al., 2014).

Previous studies of EC2 suggest its involvement in early colonization and particularly penetration success. Transient transformation of the cucurbit PM *Podosphaera xanthii* (*Pxa*) with *PxaEC2-GFP* localized it to growing hyphal tips of the fungus (Martinez-Cruz et al., 2018). This location could be consistent with a role in enhancing penetration events to form both primary and secondary haustoria as the colony expands. Overexpression of *EC2* did not alter penetration rates or spore production by adapted PM but did increase penetration success of a non-adapted PM (Schmidt et al., 2014). This suggests sufficient EC2 protein is present in compatible adapted interactions. Previous reports did not examine the impact of reducing *EC2* expression on PM growth and reproduction. Here, we show that SIGS against *GorEC2* and *EnEC2* effectively reduces spore production, by 60% and 47% respectively, in the *Gor*MGH1*-Arabidopsis* and *EnC-grapevine* systems (**Figs. 5-6**). Further analysis is needed to define the specific phase(s) of infection impacted by *EC2* reduction. Importantly, our results show that PM effectors, even those that are very highly expressed, can be targeted via SIGS to reduce PM growth and development.

### Application of SIGS against powdery mildew in agriculture and ornamental horticulture

Here, we translated our findings using the *G. orontii-Arabidopsis* system to the commercially important study of *En* infection of grapevine where new powdery mildew control methods are greatly needed. We identified nine PM targets that resulted in strongly reduced *Gor* PM proliferation on plants (**Fig. 5**). Six of these were also tested in the *E. necator*-grapevine system and SIGS against all six targets dramatically reduced *En* spore production **(Fig. 6**). This suggests effective conserved SIGs PM targets identified in one system can be adapted to control PMs of other economically important agricultural and ornamental plants. Optimization of SIGS to control powdery mildew in agricultural settings could include further refinement of the applied dsRNA sequence against a given target, modification of the dsRNA, the formulation and/or delivery methods (Cagliari et al., 2019). In addition, multiplexing dsRNAs targeting genes within a functional process/pathway or from differing processes/pathways could further increase efficacy as recently shown for dsRNAs targeting the agricultural pest whitefly (Jain et al., 2022). The benefits of topical RNAi in agriculture include its sequence specificity (and minimal off-target effects); rapid biodegradability in the environment; and little to no environmental or human health impacts (Fletcher et al., 2020). Furthermore, though regulatory requirements are not defined (Dietz-Pfeilstetter et al., 2021), topical dsRNA products are likely to be regulated as a biopesticide with potential approval extended to organic farming. Thus, topical RNAi products are likely to constitute the next wave of more sustainable methods of agricultural pest control.

## EXPERIMENTAL PROCEDURES

### Plant growth and powdery mildew inoculation

*A. thaliana* ecotype Col-0 were grown in growth chambers at 22°C, 70% relative humidity, and a 12 hr photoperiod with photosynthetically active radiation= 150µmol/(m^2^s). At 4-4.5 weeks plants were inoculated by settling tower (Reuber et al., 1998) with *G. orontii* isolate MGH1 at 2:30 pm ± 1.5 hrs. Plants for whole plant assays were grown in 4-in pots (4 plants per pot). Detached leaf assay plants were grown in plant trays (16.6 × 12.4 × 5.8-cm insert boxes; 12 plants per box; before infection, mature, fully expanded leaves were plucked and petioles inserted into 1/2x Murashige and Skoog salts/0.8% agar overlaid with Whatman 1.0, with 12-15 leaves per plate. A low to moderate inoculum of 10-14 dpi conidia (1-2 half covered leaves) was used for hyphal assays; a heavy dose (4-5 fully infected leaves) for spore counting.

*E. necator* C-strain was obtained from Andrew Walker, UC Davis, and maintained on detached leaves of Chardonnay and Carignan grapevine varieties. Grapevine hard cuttings were from Foundation Plant Services, UC Davis and grown at UC Berkeley Oxford Tract Greenhouse. Cuttings were rooted on a mist bench, transferred to 2-gallon pots with supersoil and fertilized with Osmocote. Grapevine plants were sprayed with sulfur to keep them mildew-free prior to assessment. Three leaves from the 3^rd^ or 4^th^ node of 4-month-old potted Chardonnay plants were sprayed with ∼60,000 *En* spores/mL for the whole plant assays or used in leaf disc assays inoculated with a heavy dose by settling tower.

### Powdery mildew quantification

#### Hyphal length measurements

Infected leaves or leaf discs were fixed, cleared in 3:1 (v/v) ethanol:acetic acid, and stained with calcofluor (CF, F3543, Sigma-Aldrich, St. Louis, MO) to visualize and quantify fungal growth. Cleared leaves were washed with water for 5 min, stained with 6 μg/mL CF solution for 15 min, rinsed with water for 10 min, and mounted on slides in 50% glycerol. Fungal colonies were imaged using a 10X objective of Zeiss Axio Imager fluorescence microscope, Qimaging QiClick and Micropublisher cameras, and Sutter Instruments Lambda LS Light Source with DAPI Long Pass filter set. Hyphal lengths were measured using image analysis software Fiji: ImageJ:https://imagej.net/Fiji/Downloads.

#### Spore quantification

12-15 mature, fully expanded *A. thaliana* leaves were harvested per condition at 8-10 dpi for whole plant or detached leaf assays. For grapevine, 3 leaves were collected at ∼16 dpi for whole plant assays, and 12-15 discs from leaves harvested at 14-21 dpi used for leaf disc assays. Tissue was vortexed in 15 mL 0.01% Tween-80, filtered through 40 μm mesh, and centrifuged. Spore pellets were resuspended to 200 μL to 1,200 μL with water. For each sample, nine 1 × 1 mm^2^ fields of a Neubauer-improved hemocytometer were counted and normalized to the fresh weight of the plant tissue. Mean spore count was normalized to the control for each replicate.

### RNA fluorescence labeling and uptake

Fluorescein-labeled 300 base β-actin-Mouse was synthesized using MAXIscript T7 Transcription Kit (AM1312, Invitrogen by Fisher Scientific, Waltham, MA) and Fluorescein RNA Labeling Mix (11685619910, Roche, Basel, Switzerland). To induce germination, 12 dpi spores were tapped onto a glass slide, placed in a petri dish containing water, covered in foil for 1 hr, unwrapped and left overnight in ambient light. Spores were blotted onto 1% agar pad on glass slide before applying 20 μL water or water plus Fluorescein-labeled RNA (800 ng). Spores were incubated with RNA for 90 min before imaging with a Zeiss Axio Imager fluorescence microscope. For RNase treated samples, 1 μL each of Benzonase Nuclease and RNase A was added 30 min before microscopy.

### RNA design and spray

pssRNAit (https://www.zhaolab.org/pssRNAit) was used to identify efficient siRNAs and dsRNAs and limit off-target gene silencing in the host *A. thaliana* or *V. vinifera*. Publicly available annotated genomes for *G. orontii* MGH1 (https://mycocosm.jgi.doe.gov/Golor4/Golor4.home.html; DOE Joint Genome Institute (JGI) CSP#1657 to M. Wildermuth, Project ID 1056001) and *E. necator* C-strain (Jones et al., 2014; https://mycocosm.jgi.doe.gov/Erynec1/Erynec1.home.html) were used. Template for *in vitro* transcriptions was amplified from PM-infected tissue cDNA using Phusion polymerase (M0530S/L, New England Biolabs, Ipswich, Massachusetts) and the T7 promoter (5’-TAATACGACTCACTATAGGGGG-3’) was added to both ends. dsRNA was generated using the HiScribe T7 High Yield RNA Synthesis Kit (E2040S, New England Biolabs, Ipswich, Massachusetts), purified using the Monarch RNA Cleanup Kit (T2050S/L, New England Biolabs, Ipswich, Massachusetts), reannealed by heating to 94°C and slowly cooling to room temperature over 50 min. 21 bp siRNAs were synthesized by Horizon Discovery (Lafayette, Colorado). Plants or leaf discs were infected with PM as above and then sprayed using a gravity feed airbrush with nuclease-free water (control) or RNA dissolved in nuclease-free water after infection on day 0 and on day 2, with 40 ug RNA per treatment, except for the whole plant grapevine assays in which each grape leaf was treated with 20 ug RNA.

### RNASeq and MycoCosm analyses

#### RNASeq

RNASeq data for *Gor* infection of 4-week old *A. thaliana* was downloaded from DOE JGI Data Portal, DOI: 10.46936/10.25585/60001036, JGI CSP 1657, Project ID 1290666 to M. Wildermuth. Mature, fully expanded leaves were harvested for each time point, with three independent biological replicates per time point. RNA read counts were normalized using DESeq2 package (v1.36) in R Studio v3.3.0 (Love et al., 2014).

#### MycoCosm

MycoCosm (Grigoriev et al., 2014) MCL clustering analysis of each *Gor* protein was used to assess prevalence in published PM genomes *Bgh* DH14 (Frantzeskakis et al., 2018), *Bgt* 99644 (Muller et al., 2019), *En* C-strain (Jones et al., 2014), *G. cichoracearum* UMSG1 and UMSG3 (Wu et al., 2018), and early diverging *Parauncinula polyspora* (Frantzeskakis et al., 2019), and in other published Leotimycete genomes, including those that colonize plants such as *Botrytis cinerea* (Anselem et al., 2011, Staats & van Kan, 2012), *Sclerotinia sclerotiorum* (Anselem et al., 2011), *Cadosphora sp*. SDE1049 (Knapp et al., 2018), and *Oidiodendron maius* (Kohler et al., 2015, Martino et al., 2018). Domain structure analysis including similarity by E-value and annotated enzymatic reactions by EC number were also considered. BLASTP was performed for proteins absent by MCL analysis. Reciprocal BLASTP analysis of *A. thaliana* and *Gor* proteins determined best hits based on E-values with coverage and percent identity. Annotated protein domains (e.g. Mistry et al., 2021) and EC numbers for the top *A. thaliana* and *Gor* protein hits were also examined.

## ACKNOWLEDGEMENTS

This research was supported by awards to M.C.W. from the American Vineyard Foundation, National Science Foundation MCB-1617020 and PFI-TT-1919244, and the USDA National Institute of Food and Agriculture, Hatch Project Accession Number 1016994. The work on *G. orontii* MGH1 genome sequencing, assembly, and annotation and RNASeq time course data (proposal: 10.46936/10.25585/60001036) conducted by the U.S. Department of Energy Joint Genome Institute (https://ror.org/04xm1d337), a DOE Office of Science User Facility, is supported by the Office of Science of the U.S. Department of Energy operated under Contract No. DE-AC02-05CH11231. We thank Kerrie Berry and Chris Daum and their teams for coordination of JGI data productions. For their assistance we thank: J. Jaenisch (UC Berkeley), Figure 4 diagram; S. Upadhyaya (UC Berkeley), DESeq2 analysis; A. Walker and S. Riaz (UC Davis), grapevine system setup; and K. Ryan and H. Xue (UC Berkeley) for manuscript review. A.G.M., J.T., and M.C.W are co-inventors on INHIBITORY RNA FOR THE CONTROL OF PHYTOPATHOGENS, PCT/US2022/025330, April 19, 2022; priority date April 19, 2021.

## DATA AVAILABILITY STATEMENT

*G. orontii* MGH1 (Golor4) genome is available for download at https://mycocosm.jgi.doe.gov/Golor4/Golor4.home.html from DOE JGI Data Portal, JGI CSP 1657, Project ID 1056001 to M. Wildermuth. RNASeq time course data for *G. orontii* MGH1 infection of 4-week old *A. thaliana* is available for download from DOE JGI Data Portal, DOI: 10.46936/10.25585/60001036, JGI CSP 1657, Project ID 1290666 to M. Wildermuth.

## AUTHOR CONTRIBUTIONS

A.G.M., J.T., and M.C.W. planned and designed the research, analyzed and interpreted the data, and wrote the manuscript. A.G.M., J.T., K.Y., and X.S. conducted the experimental work. K.L., S.H., V.S., and I.V.G. provided the assembled and annotated *G. orontii* MGH1 genome and RNASeq data.

## SUPPORTING INFORMATION LEGENDS

**SI Fig. 1**. *G. orontii* MGH1 gene expression shown for spray-induced gene silencing targets with predicted roles in glycogen metabolism. Mean DESeq2 normalized gene expression (± SEM) from triplicate samples harvested at 0, 6, 12, 24, 72, and 120 hours post infection (hpi).

**SI Table 1. Powdery mildew targets and primers**

