## Supplementary figures and images for "SPRAY-INDUCED GENE SILENCING IDENTIFIES PATHOGEN PROCESSES CONTRIBUTING TO POWDERY MILDEW PROLIFERATION"

### Supplemental Figure 1

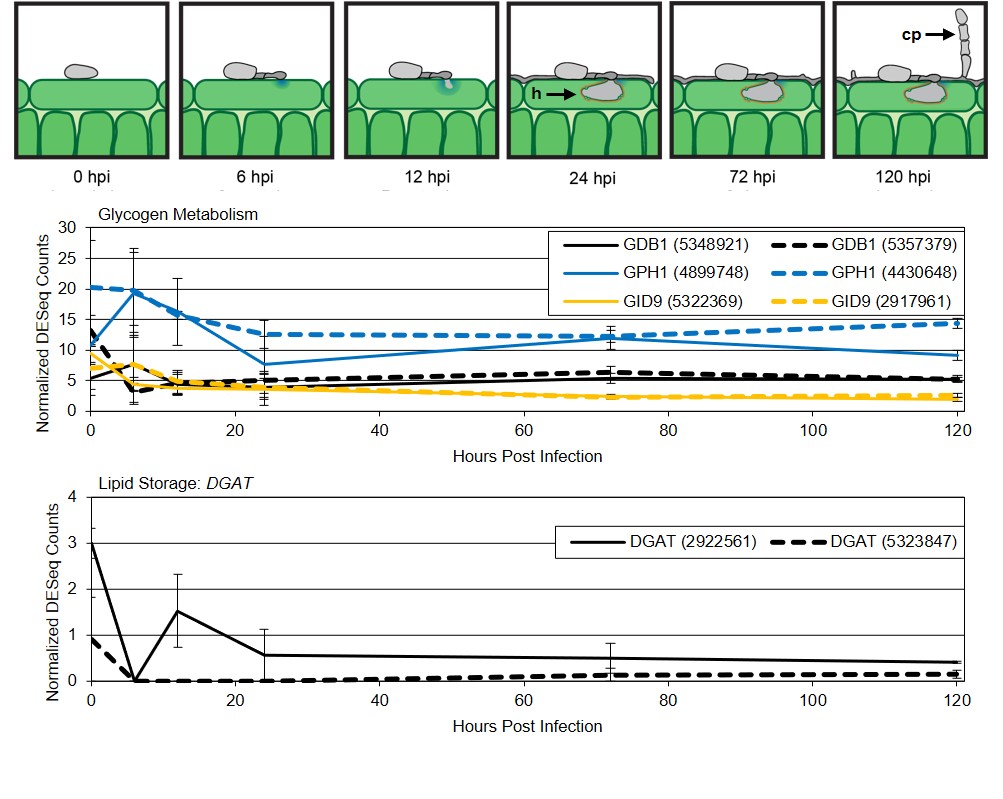
